# Symbiosis between Patescibacteria and Archaea discovered in wastewater-treating bioreactors

**DOI:** 10.1101/2022.04.10.487813

**Authors:** Kyohei Kuroda, Kyosuke Yamamoto, Ryosuke Nakai, Yuga Hirakata, Kengo Kubota, Masaru K. Nobu, Takashi Narihiro

**Affiliations:** Bioproduction Research Institute, National Institute of Advanced Industrial Science and Technology (AIST), 2-17-2-1 Tsukisamu-Higashi, Toyohira-ku, Sapporo, Hokkaido, 062-8517 Japan; Bioproduction Research Institute, National Institute of Advanced Industrial Science and Technology (AIST), Central 6, Higashi 1-1-1, Tsukuba, Ibaraki 305-8566, Japan; Department of Frontier Sciences for Advanced Environment, Graduate School of Environmental Studies, Tohoku University, 6-6-06 Aramaki Aza Aoba, Aoba-ku, Sendai, Miyagi 980-8579, Japan

**Keywords:** Candidate Phyla Radiation (CPR), Patescibacteria, Archaea, methanogenesis, symbiosis, fluorescence *in situ* hybridization (FISH), transmission electron microscopy (TEM)

## Abstract

Each prokaryotic domain, Bacteria and Archaea, contains a large and diverse group of organisms characterized with ultrasmall cell size and symbiotic lifestyles – Patescibacteria (also known as Candidate Phyla Radiation/CPR) and DPANN archaea. Cultivation-based approaches have revealed that Patesibacteria and DPANN symbiotically interact with bacterial and archaeal partners/hosts respectively, but cross-domain symbiosis/parasitism has never been observed. Here, we discovered physical interaction between ultramicrobacterial Patescibacteria and methanogenic archaea using cultures from anaerobic wastewater treatment sludge. In the cultures, we observed physical attachment of ultramicrobial cells to cells resembling *Methanothrix* and *Methanospirillum* using transmission electron microscopy and successfully detected physical association of *Ca*. Yanofskybacteria and *Methanothrix* using fluorescence *in situ* hybridization (FISH) (other ultramicrosized bacterial cells, presumably Patescibacteria, were also observed to attach on *Methanospirillum*). This was further confirmed to be a symbiosis rather than simple aggregation based on the observation that most ultramicrobacterial cells attached to *Methanothrix* were *Ca*. Yanofskybacteria and positive correlation (*p* < 0.05) between the relative abundance of Patescibacteria lineages and methanogenic archaea (*e.g., Ca*. Yanofskybacteria–*Methanothrix* and uncultured clade 32-520–*Methanospirillum*). The results shed light on a novel cross-domain symbiosis and inspire potential strategies for culturing CPR/DPANN.

## Introduction

One major lineage of the domain Bacteria, Patescibacteria (also known as Candidate Phyla Radiation/CPR) [1, 2], is a highly diverse group of bacteria, widely inhabiting [3, 4] natural and artificial ecosystems [5, 6], characterized with small cell/genome size [7] and poor (genomically predicted) abilities to synthesize cellular building blocks [4] suggestive of symbiotic dependency for cell growth. Most members remain uncultured, leaving major knowledge gaps in the range and nature of their symbioses, but several cultivation-based studies have demonstrated host-specific symbiotic/parasitic interactions between Patescibacteria and other bacteria [8–11]. This pointed towards specialization towards bacteria-bacteria symbioses, especially given parallels with DPANN, an archaeal analog who has only been observed to interact with other archaea [4, 12]; however, one study has reported Patescibacteria-eukarya symbiosis [13], indicating that the Patescibacteria host range reaches beyond bacteria. Here, based on our previous observations of positive correlations between abundances of Patescibacteria members and methanogenic archaea (“methanogens”) [6], we hypothesized that some Patescibacteria may symbiotically interact with archaea and used exogenous archaea to culture potential archaea-dependent Patescibacteria.

## Results and Discussion

To create conditions conducive for growth of archaea-dependent Patescibacteria, we took a strategy similar to virus/phage cultivation, in which exogenous methanogenic archaea are grown in the presence of Patescibacteria that will presumably grow using molecules derived from these active hosts (*i.e*., symbiosis/parasitism). We chose acetate-utilizing methanogens as partners as they inhabit methanogenic ecosystems ubiquitously [5, 6, 14], form symbiotic interactions with bacteria [14], utilize an energy source (acetate) generally non-inhibitory to organotrophs (unlike other methanogen substrates like H_2_ or formate [15]), and conveniently have highly unique cell morphology/structure easily differentiable from other organisms (*i.e*., easily traceable under the microscope). To culture Patescibacteria that may interact with archaea, we used microbial community samples from a bioreactor (“sludge”) particularly abundant in Patescibacteria as a starting material and amended them with acetate-utilizing methanogens (*Methanothrix soehngenii* GP6 and *Methanosarcina barkeri* MS) as symbiotic partners and acetate as an energy source for the archaea. Potential growth factors (yeast extract, various amino acids, and nucleoside monophosphates) were also provided as Patescibacteria are known to have poor biosynthetic capacities [4].

In the cultivation experiments, we performed serial dilutions (10^−1^ to 10^−6^) of the sludge-methanogen mixture to help eliminate low-abundance bacteria that may interfere with Patescibacteria growth. In some cultures with confirmed gas production (Table S1), we observed that many cells with morphology characteristic of *Methanothrix* (long rods with blunt ends and approximately 0.8 μm diameter) were physically and consistently associated with some ultramicrobacterial cells (< 1 μm diameter), with the number of attached cells increasing as the culture aged (Fig. S1A and S1B). To further eliminate other non-target populations in the culture (*i.e*., “enrich” the target organisms), we sub-cultured those abundant in ultramicrobacterial cells and microscopically confirmed continued physical attachment of small cells with cells with morphology characteristic of *Methanothrix* (Fig. S1C and S1D).

Using transmission electron microscopy (TEM) to observe cells in the acetate-amended enrichment culture containing the seed sludge, *Methanothrix*, and *Methanosarcina* (10^−2^ dilution) after 40 days of growth, we confirmed that coccoid-like sub-micron cells (0.46 ± 0.13 μm long and 0.36 ± 0.07 μm wide) were attached on sheathed filamentous cells highly characteristic of *Methanothrix* [16, 17] (Fig. 1A–1C). In some cases, we observed indentations of the (typically straight) cell wall of *Methanothrix* cells where the sub-micron cells were attached (Fig. 1C), suggesting that the symbiotic interaction may have a negative influence on the host cell (*e.g*., parasitism). We also found association of sub-micron cells (0.41 ± 0.05 μm long and 0.29 ± 0.05 μm wide) attached to rod-shaped cells with sheath/plug structure highly similar to *Methanospirillum* [18] (Fig. 1D), which persisted from the seed sludge. The small cells were consistently surrounded in extracellular polymeric substances (EPS)-like substances and attached to the plug structure of *Methanospirillum* cells (Fig. 1D–1F). The calculated cell volumes of the putative symbionts of *Methanothrix* (0.0377 ± 0.0200 μm^3^) and *Methanospirillum* (0.0212 ± 0.0085 μm^3^) were smaller than 0.1 μm^3^ (*i.e*., ultramicrobacterial [7]), much like Patescibacteria. Supporting the above observations, we found positive and statistically significant correlation (*p* < 0.05 based on Pearson correlation analysis) between the relative abundances of Patescibacteria and methanogenic archaea (based on 16S ribosomal RNA gene microbial community analysis in Table S2): *Ca*. Yanofskybacteria–*Methanothrix* (*r*^2^ = 0.80, Fig. S2A), uncultured linage 32-520–*Methanospirillum* (*r*^2^ = 0.70, Fig. S2B), *Ca*. Moranbacteria–*Methanolinea (r^2^* = 0.68, Fig. S2C), and *Ca*. Magasanikbacteria–*Methanolinea* (*r*^2^ = 0.91, Fig. S2C) (Table S2 and S4).

**Fig. 1.**
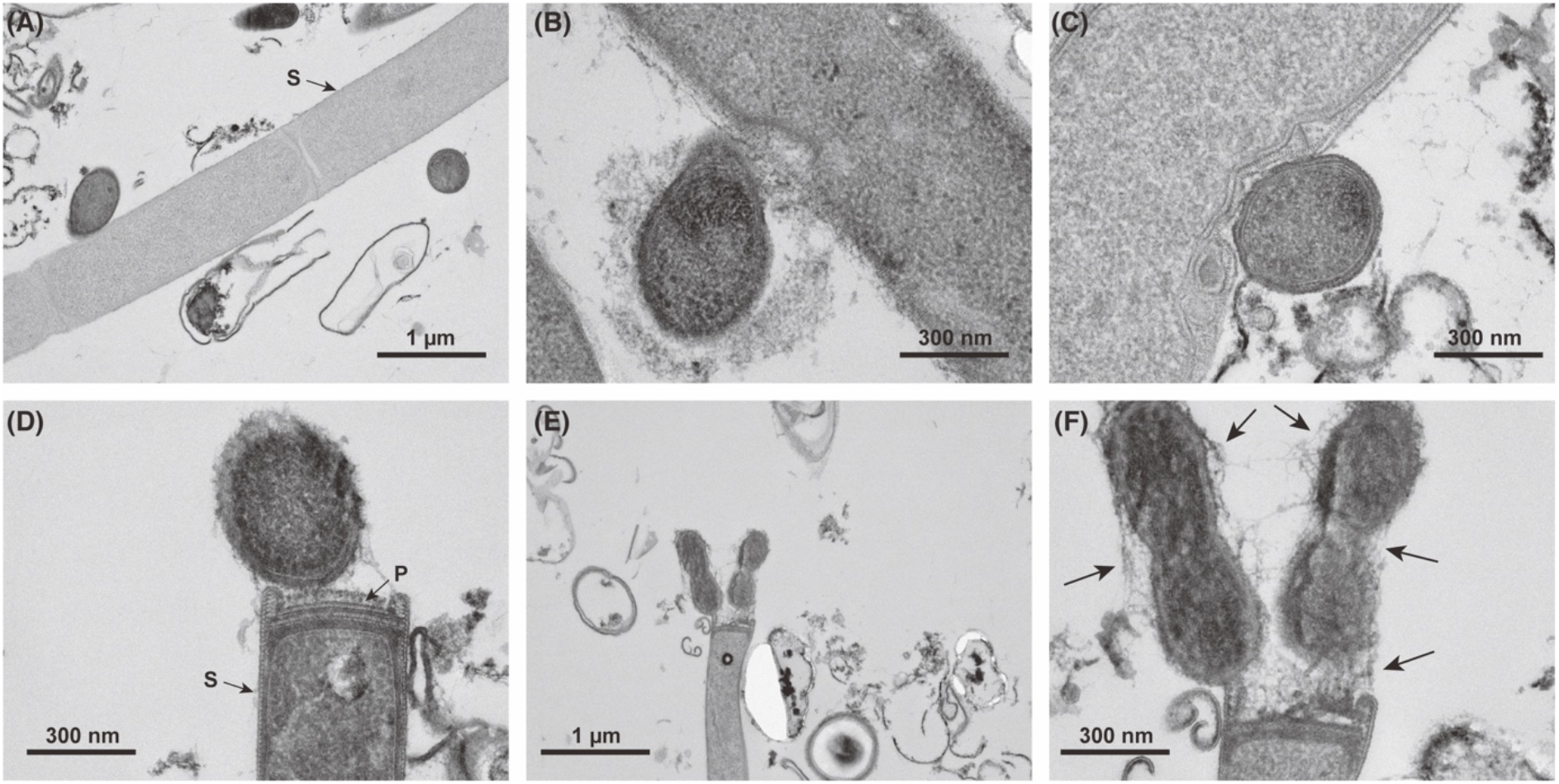
Transmission electron micrographs of small coccoid-like sub-micron cells attached on the *Methanothrix*-like cells (A–C) and *Methanospirillum*-like cells (D–F) in culture system A-d2 on days 40. P and S are plug and sheath structures, respectively. Black arrows in (F) indicate extracellular polymeric substances (EPS)-like substances.

Through fluorescence *in situ* hybridization (FISH) that allows differentiation of target populations with fluorescence microscopy, we successfully verified that the ultramicrobacterial cells attached to the surface of *Methanothrix* were indeed *Ca*. Yanofskybacteria (with probes MX825 and Pac_683 respectively; Fig. 2C and Fig S5C). In some cases, *Ca*. Yanofskybacteria cells with ribosomal activity (as confirmed by FISH, which target ribosomal RNA) were found attached to *Methanothrix* cells with low or no ribosomal activity (*i.e*., no fluorescence) (Fig. 2D and Fig. S5D), further supporting the possibility that these Patescibacteria parasitize *Methanothrix*. Moreover, the observation that nearly all *Ca.* Yanofskybacteria cells in the culture associated with *Methanothrix* (*i.e*., extremely few cells detached from *Methanothrix* and no association with other methanogens and) further evidences the specificity of the interaction (*i.e*., host specificity of *Ca.* Yanofskybacteria). We also confirmed that the ultramicrobacterial cells attached to *Methanospirillum* (cells that fluoresce with *Methanomicrobiales*-specific MG1200 but not *Methnaolinea-specific* NOBI633) were bacteria (Fig. S6), though we could not determine their phylogenetic identity. As also observed for *Methanothrix*, symbiont-associated *Methanospirillum* cells had low ribosomal activity/fluorescence (Fig. S6C), perhaps pointing towards parasitism.

**Fig. 2.**
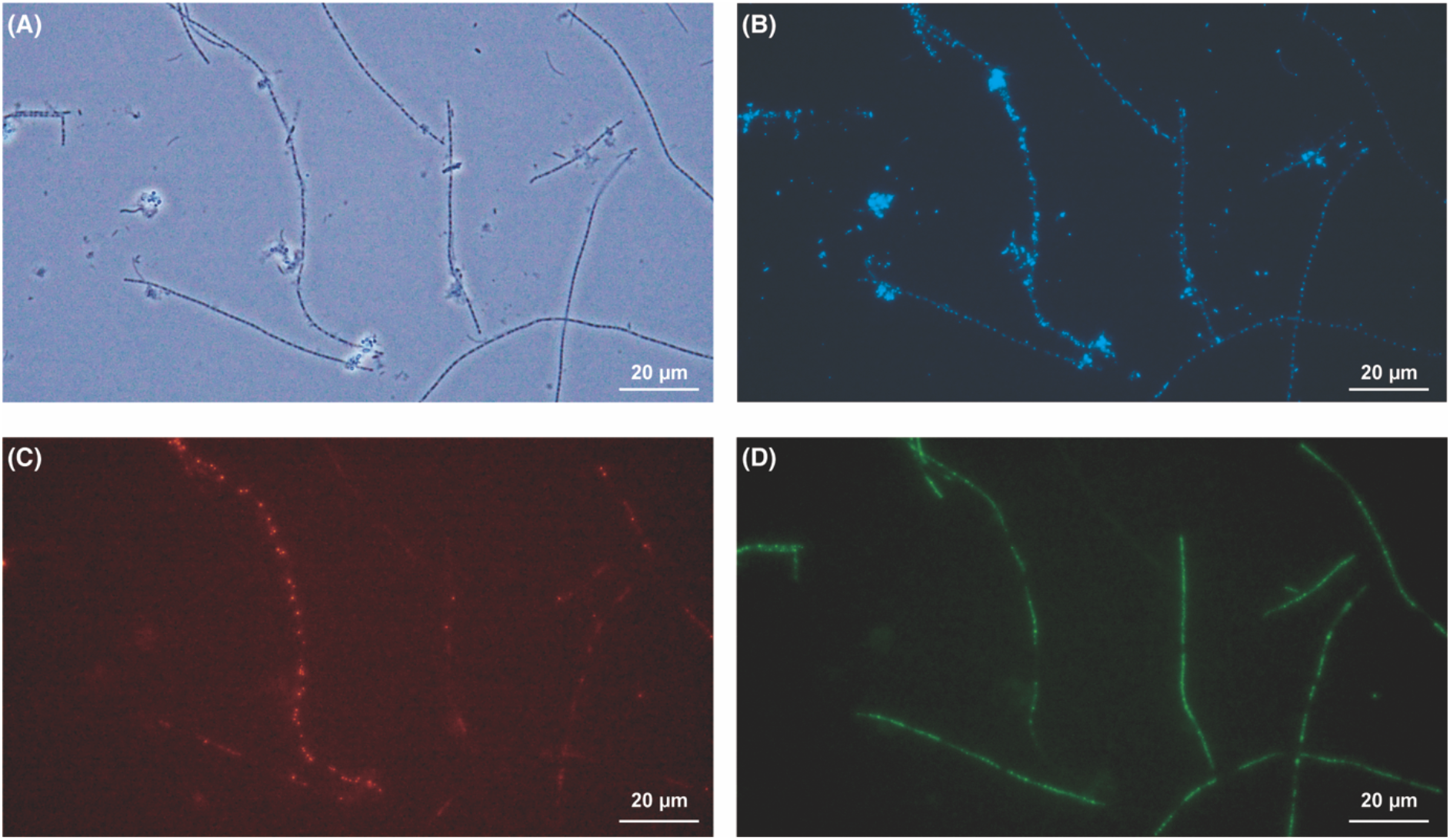
Micrographs of (A) Phase-contrast, (B) 4’,6-diamidino-2-phenylindole dihydrochloride staining, and (C) and (D) Fluorescence *in situ* hybridization obtained from culture system B-d1-d1 on days 23. (C) *Ca.* Yanofskybacteria-targeting Pac_683-Cy3 probe and (D) *Methanothrix-targeting* MX825-FITC probe.

In total, through the first successful cultivation/enrichment of the Patescibacteria class *Ca*. Paceibacteria (which *Ca.* Yanofskybacteria belongs to), we discovered that Patescibacteria/CPR can symbiotically interact with the domain *Archaea* and the obtained results suggested that the observed *Ca*. Yanofskybacteria–*Methanothrix* interaction is likely a host-specific symbiosis with negative influence on the host (*i.e.*, parasitic). The ability of Patesibacteria to interact with methanogenic archaea, a central group of organisms in anerobic ecosystems, potentially has major implications for both ecology and carbon cycling. As the presented archaea co-cultivation strategy was effective in culturing Patescibacteria inhabiting methanogenic environments, we anticipate that further refinement of co-cultivation combined with gene/protein expression will allow characterization of the details of Patescibacteria-archaea symbioses (*e.g*., parasitic or mutualistic), determination of the diversity of archaea-dependent Patescibacteria, and, ultimately, elucidation of these organisms’/interactions’ influence on anaerobic ecology.

## Acknowledgments

This study was partly supported by the Japan Society for the Promotion of Science (JSPS) KAKENHI Grant numbers 16H07403 and 21H01471, a matching foundation between National Institute of Advanced Industrial Science and Technology (AIST) and Tohoku University, and research grants from the Institute for Fermentation, Osaka (Grant No. G-2019-1-052 and G-2022-1-014). The authors thank Riho Tokizawa, Yuki Ebara, and Tomoya Ikarashi at the AIST for their technical assistance.

## Contributions

K. Kuroda and T.N. designed this study. K. Kuroda performed bioreactor sludge sampling, cultivation, microscopic observation, and 16S rRNA gene amplicon sequence analysis. K. Kuroda, K.Y., R.N., M.K.N., K. Kubota, and T.N. conducted data interpretation. K. Kuroda, M.K.N., and T.N. wrote the manuscript with input from all co-authors. All authors have read and approved the manuscript submission.

