## Supplementary figures for "Symbiosis between Patescibacteria and Archaea discovered in wastewater-treating bioreactors"

Kyohei Kuroda[1*], Kyosuke Yamamoto[1], Ryosuke Nakai[1], Yuga Hirakata[2], Kengo Kubota[3], Masaru K. Nobu[2*], and Takashi Narihiro[1*]

[1]Bioproduction Research Institute, National Institute of Advanced Industrial Science and Technology (AIST), 2-17-2-1 Tsukisamu-Higashi, Toyohira-ku, Sapporo, Hokkaido, 062-8517 Japan
[2]Bioproduction Research Institute, National Institute of Advanced Industrial Science and Technology (AIST), Central 6, Higashi 1-1-1, Tsukuba, Ibaraki 305-8566, Japan
[3]Department of Frontier Sciences for Advanced Environment, Graduate School of Environmental Studies, Tohoku University, 6-6-06 Aramaki Aza Aoba, Aoba-ku, Sendai, Miyagi 980-8579, Japan

[*]Co-corresponding authors:
Takashi Narihiro,
Masaru K. Nobu,
Kyohei Kuroda,

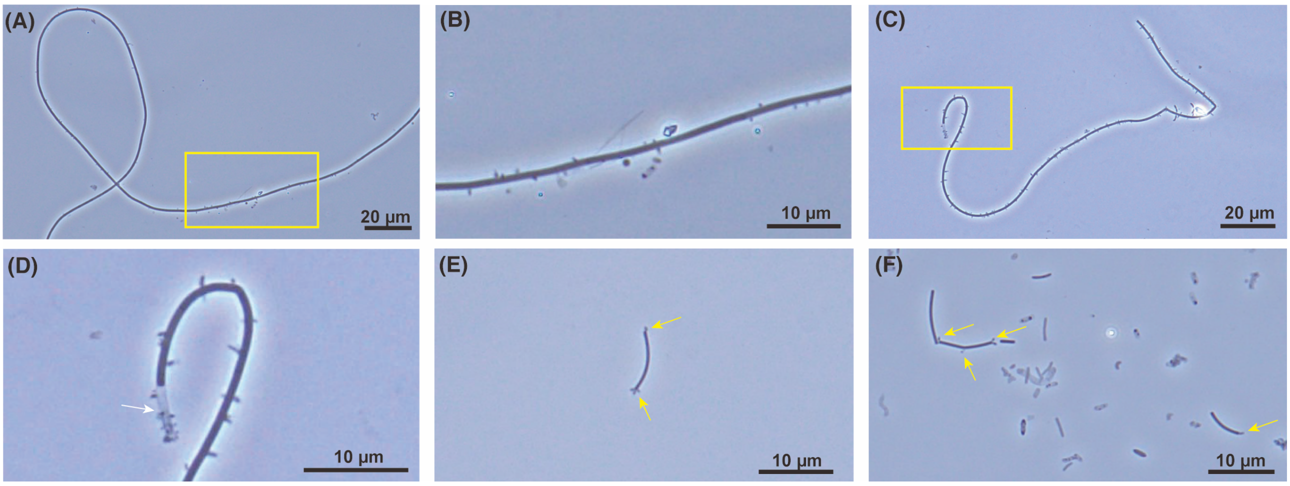


**Fig. S1.** Phase-contrast micrographs of (A)–(D) the small cells attached on *Methanothrix*-like cells in culture systems A-d2 on days 33 (A and B) and C-d2-d1 on days 23 (C and D). (E) and (F): the small cells attached on rod-shaped cells in culture system A-d2 on days 12 and 23, respectively. Yellow arrows indicate attached microbial cells on the rod-shaped cells. Yellow squares indicate high magnification parts of (A) and (B). White arrows indicate colorless cells of *Methanothrix*-like structure.

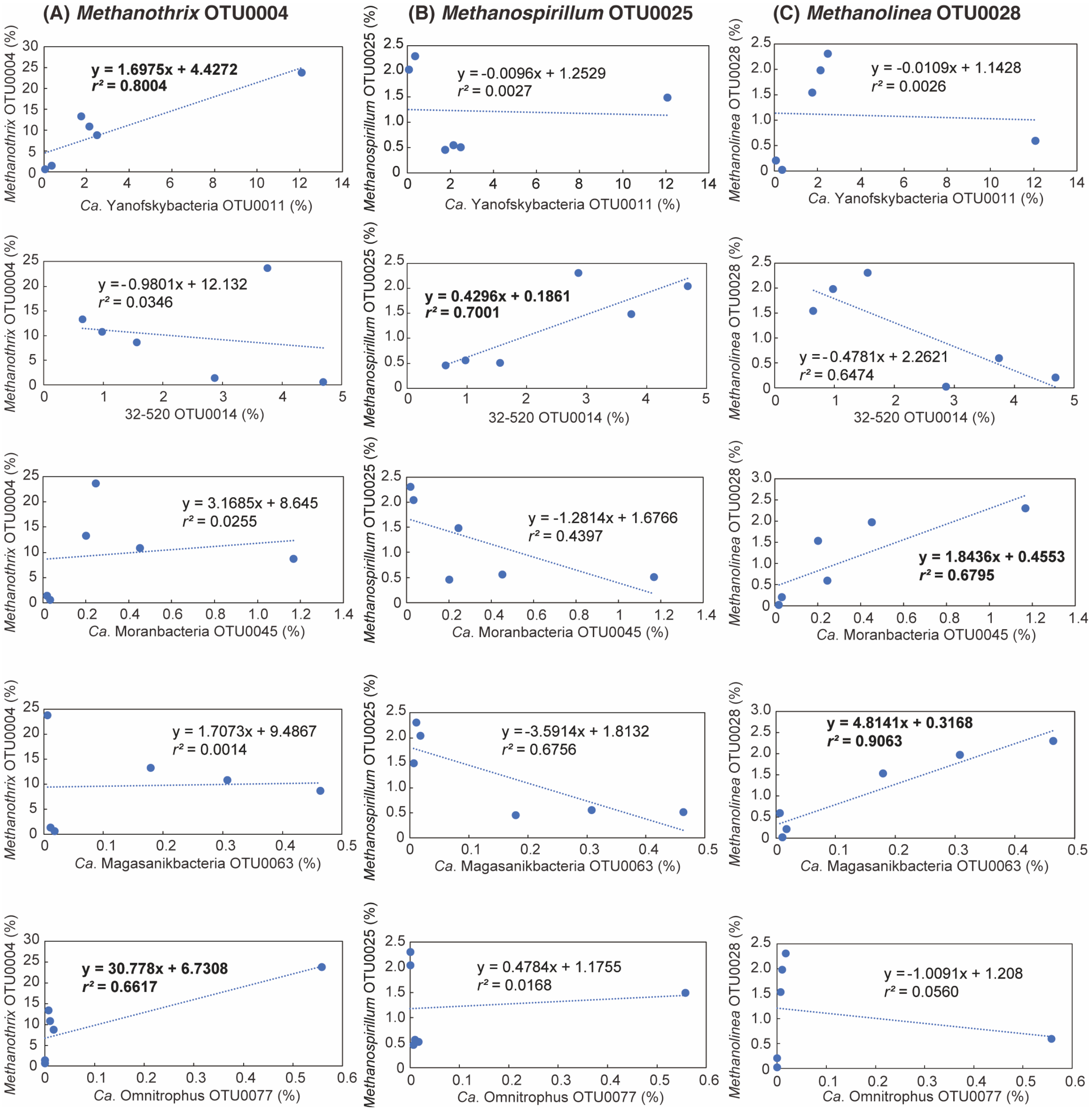


**Fig. S2.** Linear regression analysis between predominant methanogenic archaea and Patescibacteria/Omnitrophota based on 16S rRNA gene-based relative abundance. (A) *Methanothrix* OTU0004, (B) *Methanospirillum* OTU0025, and (C) *Methanolinea* OTU0028 were chosen for the regression analysis.

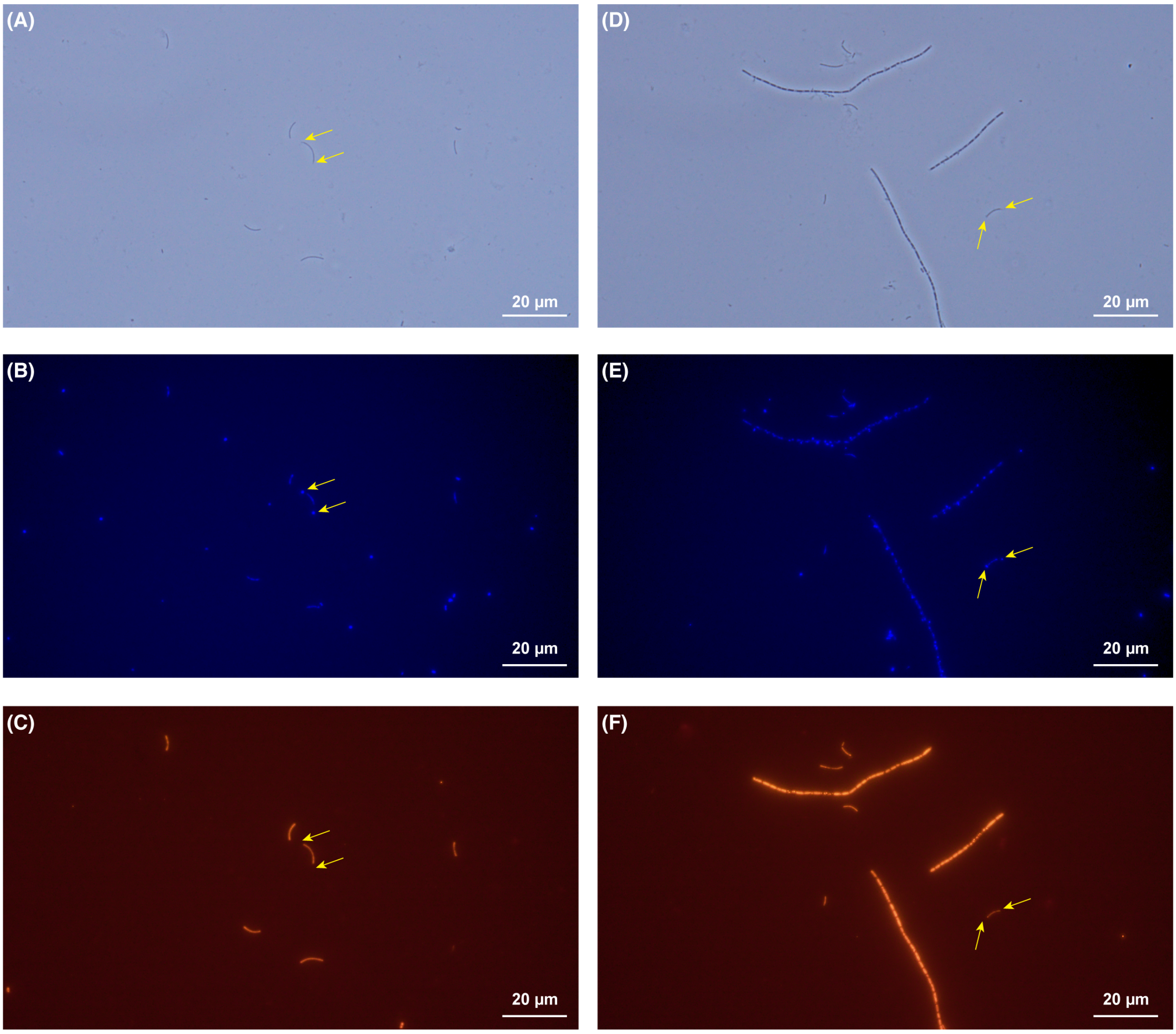


**Fig. S3.** (A) and (D) Phase-contrast, (B) and (E) 4',6-diamidino-2-phenylindole dihydrochloride staining, and (C) and (F) Fluorescence *in situ* hybridization micrographs targeting domain Archaea by ARC915-Cy3 probe obtained from culture system A-d2-d1 on days 23. Yellow arrows indicate attached microbial cells on the rod-shaped cells.

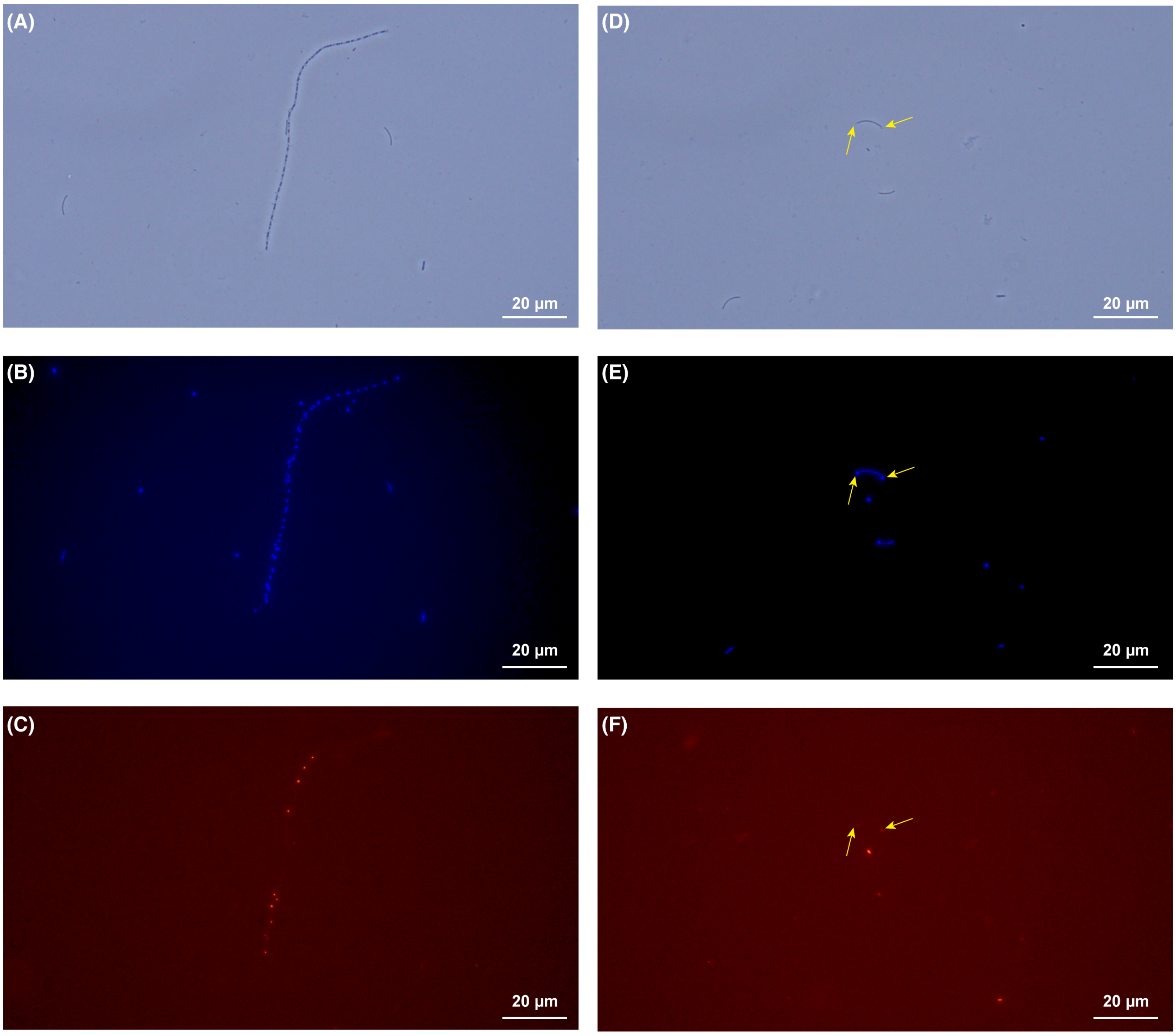


**Fig. S4.** (A) and (D) Phase-contrast, (B) and (E) 4',6-diamidino-2-phenylindole dihydrochloride staining, and (C) and (F) Fluorescence *in situ* hybridization micrographs targeting domain Bacteria by EUB338mix-Cy3 probe obtained from culture system A-d2-d1 on days 23. Yellow arrows indicate attached microbial cells on the rod-shaped cells.

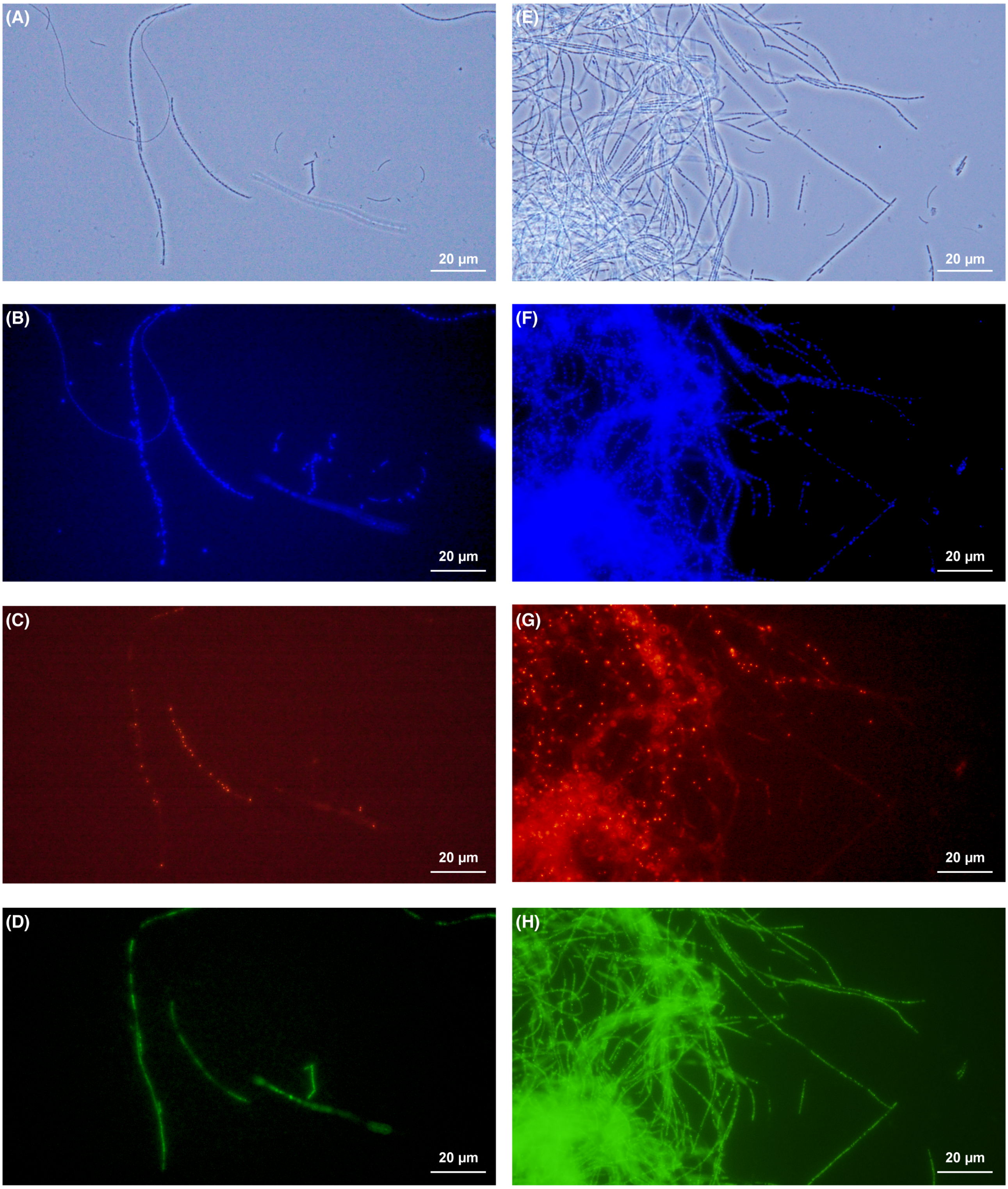


**Fig. S5.** (A) and (E) Phase-contrast, (B) and (F) 4',6-diamidino-2-phenylindole dihydrochloride staining, (C), (D), (G), and (F) Fluorescence in situ hybridization micrographs of obtained from culture system A-d2-d1 on days 23. (C) and (G) UBA9983 (Paceibacteria)-targeting Pac_683-Cy3 probe and (D) and (H) *Methanothrix*-targeting MX825-FITC probe.

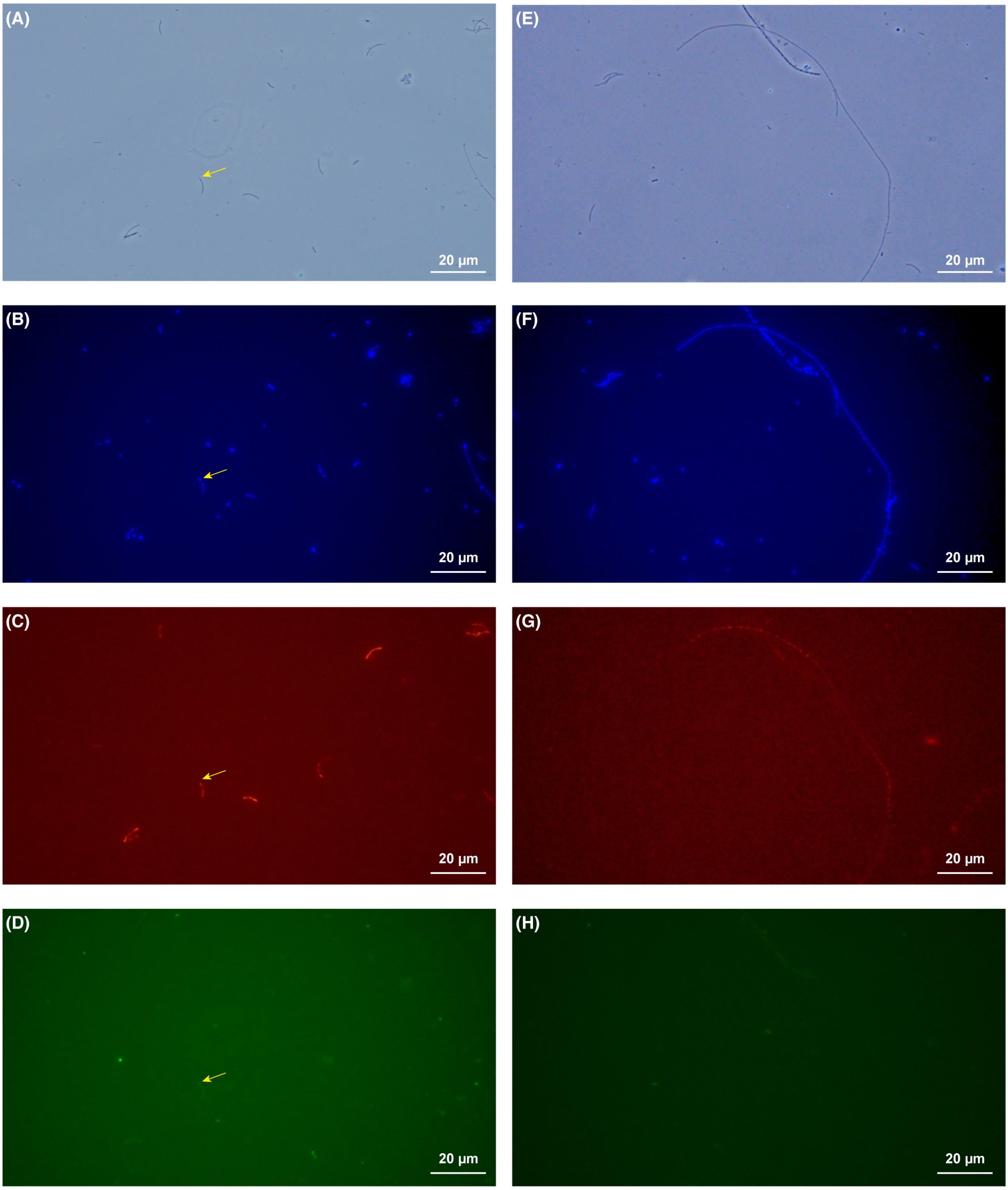


**Fig. S6.** (A) and (E) Phase-contrast, (B) and (F) 4',6-diamidino-2-phenylindole dihydrochloride staining, (C), (D), (G), and (F) Fluorescence in situ hybridization micrographs of obtained from culture system A-d2-d1 (A–D) and B-d1-d1 (E–H) on days 23. (C) Methanomicrobiales-targeting MG1200-Cy3 probe, (D) and (H) domain Bacteria-targeting EUB338mix-FITC probe, and (G) *Methanolinea*-targeting NOBI633-Cy3 probe. Yellow arrows indicate attached bacterial cells on the *Methanospirillum* cells.

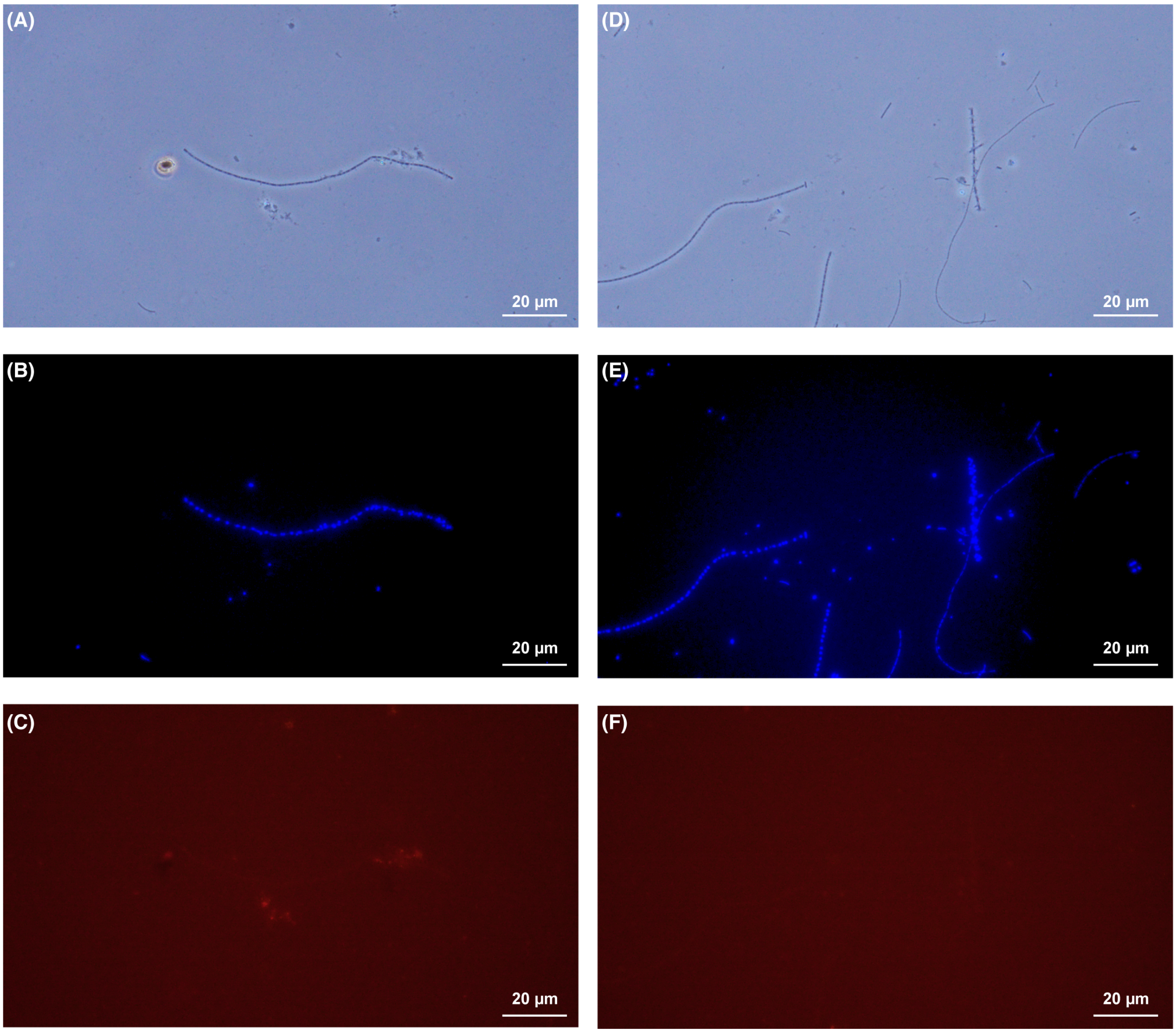


**Fig. S7.** (A) and (D) Phase-contrast, (B) and (E) 4',6-diamidino-2-phenylindole dihydrochloride staining, and (C) and (F) Fluorescence *in situ* hybridization micrographs targeting OP3 by OP3-565-Cy3 probe obtained from culture system B-d1-d1 (A–C) and C-d2-d1 (D–F) on days 23.
