## Supporting Information for "Symbiosis between Patescibacteria and Archaea discovered in wastewater-treating bioreactors"

Kyohei Kuroda[1*], Kyosuke Yamamoto[1], Ryosuke Nakai[1], Yuga Hirakata[2], Kengo Kubota[3], Masaru K. Nobu[2*], and Takashi Narihiro[1*]

[1]Bioproduction Research Institute, National Institute of Advanced Industrial Science and Technology (AIST), 2-17-2-1 Tsukisamu-Higashi, Toyohira-ku, Sapporo, Hokkaido, 062-8517 Japan
[2]Bioproduction Research Institute, National Institute of Advanced Industrial Science and Technology (AIST), Central 6, Higashi 1-1-1, Tsukuba, Ibaraki 305-8566, Japan
[3]Department of Frontier Sciences for Advanced Environment, Graduate School of Environmental Studies, Tohoku University, 6-6-06 Aramaki Aza Aoba, Aoba-ku, Sendai, Miyagi 980-8579, Japan

[*]Co-corresponding authors:
Takashi Narihiro,
Masaru K. Nobu,
Kyohei Kuroda,

### Materials and Methods

#### Enrichment culture experiments

Granular sludge collected from a lab-scale UASB reactor treating PET manufacturing synthetic wastewater were used as a microbial source for the enrichment cultures. The basal media were prepared according to previous study [1]. The enrichment culture experiments were performed at 37ºC using 50 mL of serum vials containing 20 mL medium under $N_2/CO_2$ (80:20, v/v) atmosphere. Substrates for the enrichment cultures were prepared according to Table S5. *Methanothrix soehngenii* GP6 (DSM 3671) was pre-cultivated at 37ºC for 4 weeks using the 60 mM acetate and 10 mM potassium bicarbonate as substrates. *Methanosarcina barkeri* MS (DSM 800) was pre-cultivated at 37ºC for 1 week using 10 mM methanol, 0.03% yeast extract (w/v), and 10 mM acetate as substrates. These pure cultures were transferred into the enrichment culture systems as expected host of Patescibacteria. For the enrichment cultures, we prepared 4th serial dilution ($10^{-1}$, $10^{-3}$, $10^{-4}$, and $10^{-6}$) for each culture system. Sixteen culture systems were routinely monitored by microscope equipped with phase-contrast apparatus (BX-53, Olympus, Japan) and measurement of biogas production volumes. To further enrich the targeting microorganisms, we transferred 2 mL of 1st dilution (defined as d1) of culture system B (defined as B-d1) and 2nd dilution of culture systems A (A-d2) and 3 (C-d2) into fresh medium (defined as A-d2-d1, B-d1-d1, and C-d2-d1) on days 33 under same substrates condition as shown in Table S5. The biogas compositions ($CH_4$, $CO_2$, $N_2$, and $H_2$) were determined by gas chromatography (Shimadzu, GC-8A, Japan) with a thermal conductivity detector fitted with a SHINCARBON-ST 50/80 stainless steel Column 4.0 m × 3.0 mm (ID) according to previous study [2].

Table S5. Substrates used in this study for the enrichment culture systems.

| Substrates | Concentration | Culture systems | | | |
|---|---|---|---|---|---|
| | | A | B | C | D |
| Acetate | 1 mM | Y[c] | N[d] | Y | N |
| Yeast extract | 0.03% (w/v) | Y | Y | N | N |
| Adenosine 5'-monophosphate | 0.1 mM | Y | Y | Y | Y |
| Uridine 5'-monophosphate | 0.1 mM | Y | Y | Y | Y |
| Guanosine 5'-monophosphate | 0.1 mM | Y | Y | Y | Y |
| Cytidine 5’-monophosphate | 0.1 mM | Y | Y | Y | Y |
| MEM Non-essential Amino Acids Solution (x100)[a] | 1% (w/w) | Y | Y | Y | Y |
| MEM Essential Amino Acids Solution (x50)[b] | 1% (w/w) | Y | Y | Y | Y |
| *Methanothrix soehngenii* GP6 (DSM 3671) | 0.2 mL/20 mL | Y | Y | Y | Y |
| *Methanosarcina barkeri* MS (DSM 800) | 0.2 mL/20 mL | Y | Y | Y | Y |

[a]cat no. 139-15651, FUJIFILM Wako Pure Chemical Co. Ltd., Tokyo, Japan

[b]cat no. 132-15641, FUJIFILM Wako Pure Chemical Co. Ltd., Tokyo, Japan

[c]Presence of the substrate in the medium.

[d]Absence of the substrate in the medium.

#### 16S rRNA gene sequence analysis

One mL of cultivated microorganisms was collected from 1st-batch of each enrichment culture system on days 12 (1st and 2nd dilution of 1st batch culture systems A–D) and 33 (only A-d2) and collected microbial cells by centrifugation at 17,750 g. DNA was extracted from microbial cells using FastDNA Spin Kit for Soil (MP Biomedicals, Santa Ana, California, USA) according to the manufacturer’s protocol. The 16S rRNA genes were amplified using Univ515F–Univ909R according to a previous study [3]. The PCR products were purified using a QIAquick PCR purification kit (Qiagen, Valencia, CA, USA) according to the manufacturer's protocol. The sequence analysis was performed by the MiSeq Reagent kit v3 and MiSeq system (Illumina, San Diego, CA, USA). Raw 16S rRNA gene sequences were analyzed using the QIIME 2 ver. 2021.4 [4] according to previous study [2]. The 16S rRNA gene sequences were clustered by ≥ 97% similarity to operational taxonomic units (OTUs) using vsearch software [5]. Taxonomic classification was carried out using the classify-sklearn retained on the SILVA database version 138 [6]. Pearson's correlation analysis of predominant OTUs (> 0.1% average relative abundance) was performed using the R software package ver. 4.0.2 [7].

**Fluorescence *in situ* hybridization**

On cultivation days 23, approximately 1 mL of cultivated media from 2nd-batch of enrichment culture systems A-d2-d1, B-d1-d1, and C-d2-d1 was sampled and fixed with 4% paraformaldehyde in phosphate-buffered saline (PBS) for 4 hours at 4ºC and stored in 50% ethanol with PBS at -20ºC. Fluorescence *in situ* hybridization (FISH) was performed as described previously [8]. The hybridization and washing slides were 46ºC for 6–8 hours and 48ºC for 20 mins, respectively. The FISH probes were shown in Table S6. An equimolar mixture (defined as EUB338mix) of EUB338 [9], EUB338I, EUB338II, and EUB338III [10] were used for detection of all bacteria. Formamide concentrations used in this study were follows: EUB338mix, MG1200 [11], and NOBI633 [12], 10%; ARC915 [11], 35%; MX825 [11], 20%; OP3-565 [13], 30%; and Pac_683 [14], 25%. The PAC_683 shows perfect match for predominant order *Ca.* Yanofskybacteria OTU0011 (UBA9983) and its related clones (LC177595.1, JX100399.1, and EF602504.1). All probes were labeled with FITC or Cy3. The FISH samples were also stained with 4',6-diamidino-2-phenylindole dihydrochloride (DAPI). The microscopic images were observed by epifluorescence microscope (BX-53, Olympus, Japan) with a color CCD camera (DP-74, Olympus, Japan). Phase-contrast, FISH, and DAPI micrographic images were uniformly processed across the entire images using Adjust color of Preview application on Mac OS 11.6.5 and Photoshop CC (Adobe Creative Cloud).

**Table S6**. Fluorescently labeled 16S rRNA-targeted oligonucleotide probes used in this study.

| Probes | Target group | Probe sequence (5' to 3') | Reference |
|---|---|---|---|
| EUB338 | Bacteria | GCTGCCTCCCGTAGGAGT | [9] |
| EUB338I | Bacteria | GCAGCCTCCCGTAGGAGT | [10] |
| EUB338II | Bacteria | GCAGCCACCCGTAGGTGT | [10] |
| EUB338III | Bacteria | GCTGCCACCCGTAGGTGT | [10] |
| ARC915 | Archaea | GTGCTCCCCCGCCAATTCCT | [11] |
| MX825 | *Methanothrix* | TCGCACCGTGGCCGACACCTAGC | [11] |
| MG1200 | Methanomicrobiales | CGGATAATTCGGGGCATGCTG | [11] |
| NOBI633 | *Methanolinea* | GATTGCCAGTTTCTCCTG | [12] |
| OP3-565 | *Candidatus* Velamenicoccus | TACCTGCCCTTTACACCC | [13] |
| Pac_683 | Order *Ca.* Yanofskybacteria | TCAACGGATTTCACCCCTACAC | [14] |

**Transmission electron microscopy**

On cultivation days 40, approximately 1 mL of cultivated media from culture system A-d2 was sampled and sandwiched with the copper disks in liquid propane at -175ºC. After frozen of the samples, the liquid was freeze substituted with 2% glutaraldehyde, 1% tannic acid in ethanol, and 2% distilled water at -80ºC for 2 days. Dehydration was performed by anhydrous ethanol at 3 times for 30 mins each. Infiltration was carried out by propylene oxide at 2 times for 30 mins each, and the sample was put into a ratio of 7:3 mixtures of propylene oxide and resin for 1 hours. After volatilization of propylene oxide, the sample was transferred to a fresh 100% resin, and polymerized at 60ºC for 48 hours. Ultra-thin sections at 70 nm with a diamond knife using an ultramicrotome (Ultracut UCT, Leica, Vienna, Austria). The sections were stained with 2% uranyl acetate at room temperature for 15 mins and washed with distilled water followed by secondary-stained with Lead stain solutions (Sigma-Aldrich Co., Tokyo, Japan) at room temperature for 3 mins. The grids were observation by a transmission electron microscope (TEM) (JEM-1500Plus, JEOL Ltd., Tokyo, Japan) at 100 kV. Digital images were observed with a CCD camera (EM-14830RUBY2, JEOL Ltd., Tokyo, Japan). The cell diameters were measured from 8 and 5 single-cells attached on *Methanothrix*- and *Methanospirillum*-like cells, respectively. The cell volumes were calculated according to Van Wambeke and Bianchi (1985) [15].

**Deposition of DNA sequence data**

The raw sequence data were deposited into the DDBJ Sequence Read Archive database (DRA013834). The 16S rRNA gene sequences of representative OTUs were described in Table S3.
